# Intragroup sociality drives individual participation in intergroup competition in an urban-dwelling nonhuman primate

**DOI:** 10.1101/2023.09.06.556573

**Authors:** Bidisha Chakraborty, Stefano S.K. Kaburu, Krishna N. Balasubramaniam, Pascal R. Marty, Brianne Beisner, Eliza Bliss-Moreau, Lalit Mohan, Sandeep K. Rattan, Brenda McCowan

## Abstract

There is significant variability in the extent to which individuals from group-living species participate in intergroup competition (IGC) over resources. Such variability may be particularly robust in human-impacted environments, wherein anthropogenic resources may strongly impact intergroup overlap and competition. Examining the socio-ecological causal factors driving these differences can shed new lights on animals’ costs-benefits trade-offs driving engagement in such interactions. Here we use (peri)urban-dwelling rhesus macaques (*Macaca mulatta*) as model-systems to examine how individual differences in socio-ecological characteristics (i.e., animals’ sex, rank, and affiliative connections) may influence propensities to participate in IGC in anthropogenic environments. We found that participation was strongly influenced by individuals’ affiliative connections. Interestingly, monkeys with more coalitionary support partners had higher IGC participation, but this was more pronounced for individuals that are more peripheral in their proximity and multilayer affiliative networks, compared to more socially central individuals. Moreover, males and low-ranking individuals were more likely to participate in IGC. Our findings suggest that differential access to socio-ecological resources likely drives IGC participation. The evaluation of such inter-individual differences in socio-ecological flexibility and decision-making in dynamic environments is critical for future research related to the links between behavioural ecology, health outcomes, and human-wildlife co-existence in anthropogenic landscapes.

## INTRODUCTION

Competition between groups or intergroup competition (hereafter, IGC) is a critical aspect of humans’, and almost all group-living animals’, socioecology. In animal groups, IGC can manifest as direct physical aggression between single individuals of different animal groups, between sub-groups or entire groups facing off [1], whereas, in humans, it can be in the form of modern [2,3] and primitive warfare [4,5], cattle raids in pastoral societies [6], inter-neighbourhood conflict as well as gang violence [7]. At the core of such diverse expressions of IGC is access to resources, such as money, energy, food, territory, or mates [8–13]. Consequently, success in IGC, via greater access to resources, has been shown to positively influence health and fitness across a wide range of group-living animals (reviewed in [14], including social insects [15] green woodhoopoes, *Phoeniculus purpureus* [16,17]; banded mongoose, *Mungos mungo* [10]; lions, *Panthera leo* [18]; Japanese macaques, *Macaca fuscata* [19]; chimpanzees, *Pan troglodytes* [9] and humans [20]). On the other hand, IGC can also be costly, inducing significant energetic demands, injuries, and disease, that may culminate in negative impacts on individuals’ fitness and mortality [7,9,21–28]. Indeed, such profound intergroup variation in the costs-to-benefits tradeoffs of IGC is hypothesized to be a major selective force in the evolution of cooperation and group-living and socio-ecology across the animal kingdom [29,30].

Interestingly, not all group members participate in IGC with the same frequency and intensity; many vertebrate taxa display strong inter-individual differences in initiation and participation in IGC (e.g., birds [31]; canids [32]; meerkats, S*uricatta suricata* [33]; primates [22,34–40]). Such differences in turn are associated with individual-level impacts on health [41] as well as access to food, mates or social benefits [42,43]. Various socioecological models which have been proposed to understand factors influencing such individual differences in participation in IGC, and their consequences for variation in intragroup social structure, propose that IGC is driven by varying resource characteristics, which in turn affects the degree of despotism versus social tolerance among animals within groups [44–47]. Thus, socio-ecological frameworks are useful for conducting empirical research related to understanding the relationship between animals’ propensities to participate in IGC and their intragroup socio-demographic characteristics [48–50].

While there has been extensive research implementing socio-ecological frameworks with respect to understanding the effect of IGC on intragroup social structure [51–56], we know very little about the opposite – that is, how an individual’s intragroup social relationships affect participation in IGC. This is despite a growing consensus among human and animal socioecologists that the link between IGC and intragroup interactions is very much a bi-directional, adaptive relationship [6,57,58]. In human societies, there is indeed strong evidence that individual heterogeneities in intragroup sociality shape intergroup behaviour like the formation of large-scale raiding parties [6,59]. However, we do not have a clear understanding of whether/how individual’s social connectedness might shape their behaviour in IGC contexts in other animals.

To date, research on inter-individual differences in participation in IGC has focused on the effects of animals intragroup sociodemographic attributes such as sex, age and rank on IGC [1,37]. These studies have shown discrepant and often unclear patterns. For example, since individuals with higher access to resources presumably stand to gain the most during IGC, higher participation by dominant individuals is often seen, for example, in free-ranging dogs (*Canis familiaris*) [32], meerkats [60], bonnet macaques (*Macaca radiata*) [49], vervet monkeys (*Chlorocebus pygerythrus*) [34] and chacma baboons (*Papio ursinus*) [61] and white-faced capuchins (*Cebus capucinus*) [62]. However, the opposite, i.e. higher participation by subordinate animals is also seen in nonhuman primate species like rhesus macaques (*Macaca mulatta*) [63] and Japanese macaques (*Macaca fuscata*) [64], while a recent meta-analysis on 31 primate species found no effect of rank on participation in IGC [37].

Sex-based differences in participation are also common. Male and female participation are expected to be driven by different motivations, with males participating for access to mates, and females for access to food [36,65]. Across the animal kingdom, males generally have higher rates of participation in IGC, as seen in feral horses (*Equus caballus*) [66], grey wolves (*Canis lupus*) [67], Alaotran gentle lemurs (*Hapalemur griseus alaotrensis*) [68], white-faced capuchins [69], bonnet macaques [49], Japanese macaques [64], Assamese macaques (*Macaca assamensis*) [70] and humans [71]; but similar or higher participation by females is also seen in some cases (e.g., spotted hyena, *Crocuta crocuta* [72]; ring-tailed lemurs, *lemur catta* [38]; Tibetan macaques, *Macaca tibethana*; reviewed in [73]).

The discrepant patterns of participation could be reconciled by examining individual decision-making as a culmination of the relative effects of both its intrinsic socio-demographic identity (e.g., sex, age, dominance rank) as well as the extrinsic socio-ecological factors (e.g., access to resources, their social connectedness or support). Very little is known about how animals’ intragroup social connectedness or position affects their propensities to take risks in IGC. Individual social position or social connectedness in a group can affect their access to food resources as well as social support in the context of intra-group conflicts [74–76]. Despite the clear importance of social support during IGC, the relationship between the two has rarely been explored. Few exceptional studies have indeed found evidence of favourable proximate social environment, such as presence of closely-bonded individuals, positively affecting animals’ decisions to participate in IGC [34,77–79]. However, the differential effect (if any) of various levels of individual social position on participation in IGC has not been studied in nonhuman animals. Doing so can help pinpoint the effects of individual-level heterogeneities, and specifically their social roles across multiple (rather than just single) social contexts, and their effects on their propensities to take risks.

To date, socio-ecological frameworks to understand the causal factors and consequences of IGC have largely focused on wildlife populations in natural landscapes or those with little human impact. In comparison, there is limited knowledge of the socio-ecology of IGC among wildlife in anthropogenically-impacted landscapes. Such assessments of IGC on (peri)urban wildlife populations are nonetheless important given the growing recognition that humans have been impacting wildlife ecology throughout our shared evolutionary histories [80], leading to the proposition of extending socio-ecological frameworks to include anthropogenic factors to understand the ecology and evolution of urban wildlife [81–84]. In this context, IGC may occur frequently among wildlife in anthropogenic landscapes since these are typically characterized by high population densities of wildlife (and people) being exposed to varying resource distribution and abundance [85,86]. Yet while a growing body of research is uncovering the effects of anthropogenic factors on intragroup social behaviour [87–91], their impact on intergroup behaviour remains less understood. Some exceptional studies in this regard have revealed that urbanization modifies territorial behaviour and increases the frequency of encounter rates and IGC among animals in human-modified landscapes, in species like song sparrows (*Melospiza melodia*) [31], banded mongoose [85], yellow mongoose (*Cynictis penicillate*) [92], racoons (*Procyon lotor*) [86], dogs [32,93], rhesus macaques (*Macaca mulatta*) [94], northern pig-tailed macaques (*Macaca leonina*) [95] and others [96]. However, individual-level differences in participation in IGC among urban/peri-urban environments remain virtually unknown. The few studies exploring this have been conducted on semi-provisioned groups of macaques, and have indeed found inter-individual differences in participation in IGC with higher participation by both males (bonnet macaques[49], rhesus macaques [94]) and females (Tibetan macaques [97]), as well as support for both higher [94] and lower [49] rates of IGC in groups inhabiting more anthropogenic areas. More thorough assessments are vital to understand individual motivations that drive animal behavioural ecology in human-impacted areas, and in turn, how animals’ realization of costs-benefits tradeoffs in such environments may impact their health and fitness.

Social network analysis (SNA) has provided valuable insights into individual strategies driving engagement in cooperation and IGC in humans. For example, SNA has helped uncover the importance of social centrality over dominance or kin relationships in successfully inciting livestock raids in agro-pastoralist societies [6], the social diffusion of murders in inter-gang interactions [7], as well as the patterns of reciprocity and evolution of cooperative tendencies in modern hunter-gatherer societies [98]. Nevertheless, the potential of SNA is yet to be utilized to explore individual participation in animal IGC. Moreover, an individual’s social position in a group is a result of relationships across various contexts (such as aggression or affiliation, or even across various types of affiliative interactions). Per this ‘multi-layer network’ framework, an individual that is well-connected within and across multiple types of affiliative behavioural networks compared to a single kind might have better access to social support, thus reducing their costs of participating in IGC. Thus, multilayer social networks offer an even more promising advancement in the context of understanding the drivers of IGC compared to traditional SNA approaches [99,100]. By capturing animals’ affiliative ties and relationships across various, relevant contexts, multilayer social networks may be used to understand whether complex aspects of intragroup sociality, such as an individual’s multidimensionality and importance across (in addition to within) affiliative networks, might influence their likelihood of participation in IGC.

To address these gaps in the literature, we examined individual animals’ propensities to participate in IGC, using free-ranging rhesus macaques (*Macaca mulatta*) living in anthropogenic urban landscapes as a model system. Rhesus macaques are among the most socio-ecologically flexible wildlife taxa, occupying diverse habitats from cities to agricultural fields to forested areas, wherein they feed on both natural and anthropogenic food [101]. Throughout their range, they live in multi-male multi-female social groups in which females are philopatric and males disperse [102]. There is also systematic variation in the patterning and distribution of intragroup agonistic (aggression, submissive signalling) [103,104] and affiliative (e.g., social grooming, lip-smacking, social tolerance, coalitionary support during conflicts) interactions, that in turn gives rise to profound intraspecific variation in emergent social structures [105–107]. Throughout their range, rhesus macaques regularly overlap with people, culminating in varying patterns of human-macaque interactions such as aggression and food-provisioning [108]. Yet most of our knowledge of rhesus macaque socio-ecology stems from semi free-ranging populations rather than populations within their natural range [109]. More pertinently, there exists little research on IGC in this species, with the focus largely being on demographic characteristics such as the effects of sex, rank and age [63,94,110,111] again, mainly arising from semi free-ranging populations. These studies all reported higher participation in IGC by subordinate and young males but did not delve into the socioecological complexities driving such participation decisions. Such findings also necessitate the extension of research on IGC to populations within the natural range of rhesus macaques that are also subject to varying degrees of anthropogenic impact.

We hypothesized that macaques with the greatest socio-ecological resource holding potential or capital, i.e., individuals with highest access to resources, will participate more in IGC (*top participant hypothesis*). Previous research on macaques, and indeed other nonhuman primates with similar social organizations, has revealed strong evidence for males, high-ranking individuals and socially well-connected individuals to all have greater access to resources [75,106,112–114]. We therefore predicted that these individuals will also show the greatest propensities to participate in IGC. Alternatively, individuals that have the least access to resources due to intra-group competition might be the most likely to participate in IGC to offset their needs as alternative feeding and/or intragroup social tolerance strategies (*underdog hypothesis*) [55]. Based on this hypothesis, we therefore expected females, lower-ranking individuals and socially peripheral or the least well-integrated (compared to central or the most well-integrated) individuals to show the greatest propensities to participate in IGC. Finally, since participation may prove particularly costly for individuals with the least access to resources, we expected that the participation of ‘underdog’ animals - females, subordinate individuals, and socially peripheral individuals - to especially be dependent/contingent on their access to intra-group social support to offset participation costs. Specifically, we predicted that such individuals would show greater propensities to participate in IGC if they receive more (compared to less) coalitionary support from their group members in the context of intra-group conflicts. Moreover, given that the strength of individuals’ ties across multiple (rather than just single) forms of affiliation may be especially advantageous for resource access, health and fitness [115,116], we also predicted that individuals’ centrality in their multilayer affiliative networks would have a particularly strong, positive influence on their likelihood of participation in IGC.

## ETHICAL NOTE

The protocols used in the study were approved by the Institutional Animal Care and Use Committee (IACUC) of the University of California, Davis (protocol # 20593). This study was also conducted in collaboration and consultation with the Himachal Pradesh Forest Department and adhered to the local Indian laws.

## METHODS

### Study site and subjects

This study was conducted in an urban Hindu Temple and surrounding forested areas (Jakhu Temple: 31.1008° N, 77.1845° E), in the Himalayan city of Shimla in Himachal Pradesh Northern India. It is comprised of both anthropogenic features related to the temple (e.g., paved temple grounds, few lawns and gardens, and temple administrative and residential buildings), as well as forested slopes descending from the temple to the town boundary (see [117] for more details). The site has a large population of rhesus macaques, with 5-6 groups visiting the temple regularly. It is also a popular tourist spot with hundreds of tourists visiting the temple every day, leading to frequent interactions between macaques with humans and anthropogenic landscape features. More pertinently, the intense provisioning of macaques by people, as well as the distribution and abundance of anthropogenic food such as trash bins and provisioning areas in the environment, leads to frequent IGC between macaque groups.

Data were collected on 3 groups of rhesus macaques that ranged in the study area; they were named HG, RG and SG. Subjects were 116 adult macaques from across these three groups: 23 from HG (17 females and 6 males), 36 from RG (24 females and 12 males) and 57 from SG (44 females and 13 males).

### Behavioural data collection

We collected behavioural data from July 2016 to February 2018 for 5 days a week and from 9 AM-5 PM. This followed a three-month preliminary phase in which observers identified all subjects and completed inter-observer reliability tests (details in [117]). We used HandBase application (DDH software) on Samsung Galaxy tablets, and focal animal sampling procedures [118]. Focal subjects were approached in a pre-determined, randomized order, and were followed for 10-minute periods recording data on their social and feeding behaviour. Within each focal session, we collected behavioural data of animal-animal and human-animal social interactions in a continuous manner. Specifically, we collected macaque-macaque dyadic agonistic interactions, such as aggression followed by submission, and submissive signalling behaviours such as silent bared teeth (SBT), avoidance, and fleeing. These interactions were recorded both between the focal animal and its group conspecifics, as well as between the focal animal and macaques from other groups that were both subjects (i.e., belonged to one of our study groups) and unidentified individuals. We also recorded affiliative behaviours such as grooming, huddling, and coalitionary support during an intra-group conflict. In addition to the continuous data collected as part of focal sampling, point-time samples were also collected every 2 minutes from the focal subjects, during which we recorded the identity of individuals present in body length of the focal individual. For more detailed information on behavioural data collection and the ethogram, see Supplementary Materials Table S1 and/or our previous publications [89,117,119]. In total we collected 1420.35 hours of focal sampling data across the three groups, with an average of 12.12 ±4.94 hours of observation per individual.

### Calculations of individuals’ participation in intergroup competition (IGC)

To calculate individual-level rates of participation in IGC, we calculated the frequency of direct aggressive behaviours that involved interactions between focal animals and macaques from other groups (see definitions and description above) per observation time from individuals from all 3 study groups. That is, we counted an individual’s participation in IGC during their 10-minute focal observation period as well as during other monkeys’ focal observations. This was to account for events when a non-focal individual engaged in IGC to support a focal individual (or group conspecific), or if a focal individual from one of our study groups aggressed a non-focal individual from another study group with the latter retaliating by participating. Since we quantified participation in IGC if it was observed during the focal observation period of any monkey in any of our 3 study groups, we used the combined individual observation time of individuals from all 3 study groups as an offset variable during analysis.

### Calculation of dominance ranks

Dominance rank was calculated from all dyadic interactions with clear winner-loser outcomes in aggressive interactions, using *Perc* package in R [120]. This network-based method can infer rank for sparse or missing data from non-interacting dyads, using direct as well as indirect wins- and-losses, which can offer an advantage over some other ranking methods like David’s score that is popularly used by behavioural ecologists [121–123]. We calculated male and female dominance rank separately using just female-female and male-male agonistic interactions. We did so since rhesus macaque male and female dominance rank acquisition have slightly different mechanisms with female ranks being predominantly maternally inherited and male rank being a result of maternal rank and other factors such as tenure, phenotypic quality and natal transfer status (whether they stayed with their natal group or emigrated) [114,124,125]. Moreover, to control for differences in the 3 group sizes, we standardized the dominance ranks per group to create a rank index, such that the values ranged from 0 (lowest rank) to 1 (highest rank) for each group.

### Estimations of individuals’ social support and connectedness

We calculated two measures of social/coalitionary support. First, we computed a direct measure of social support quantified as the rate *of coalitionary support received* by individuals during aggressive interactions. An individual’s higher rates of receipt of coalitionary support in intragroup contexts would presumably indicate that they would also have higher chances of being supported during IGC. Secondly, we calculated individual *eigenvector centrality of coalitionary support*. Eigenvector centrality (EC) is a social network measure that quantifies an individual’s direct connections and secondary connections, that is, the connections of those direct connections [126–129]. An individual with high coalitionary support EC is connected to many coalitionary support partners who themselves have many such partners. To calculate EC, we constructed a weighted, directed social network, in which edges were rates of coalitionary support received during the observation time in which both the individuals were concurrently present in the group. Similar to dominance rank, the EC for each group was standardized to account for different group sizes. EC of coalitionary support, grooming and social proximity (described below) was calculated using MuxViz software v2.0, which uses R environment to calculate social network measures [130,131]. MuxViz was created to calculate multilayer social network measures (discussed below), but it can also be used to calculate commonly used social network measures such as EC. For this paper, we decided to use MuxViz for all single and multilayer social network calculations so that we were able to compare the relative importance of various levels of social support. All single layer social networks were visualized using Cytoscape software v3.7.1 [132].

We also estimated individual *social position* incorporating both single as well as multiple affiliative behaviours. For the measure incorporating both direct and indirect affiliative connections in single behaviours, we calculated *eigenvector centrality of grooming* as well as *social proximity eigenvector centrality* (defined as number of individuals present within their body-length while engaging in affiliative interactions such as grooming, huddling or coalitionary support). An individual with high grooming or social proximity EC is connected to many grooming partners/individuals in proximity who themselves have a large number of such partners, thus, implying that such individuals generally occupy a central position in the group. To calculate social position incorporating multiple types of affiliation, we used a multilayer network approach [133]. Specifically, we calculated each macaque’s *multilayer versatility* (multilayer equivalent of single layer eigenvector centrality) or hereafter, ‘*affiliative versatility*’ that incorporates its direct and indirect connectedness across multiple (rather than single) affiliative networks behaviour, in this case grooming, coalitionary support, huddling, and social proximity behaviour (Figure 1). High affiliative versatility would be indicative of high *social integration*, i.e., many direct and indirect social connections within and across all affiliative behaviours. All measures of social position, i.e., EC of grooming, social proximity as well as eigenvector versatility was calculated in MuxViz software v2.0 and standardized to account for group size. All multilayer visualizations were also created using MuxViz graphical user interface v2.0.

**Figure 1:**
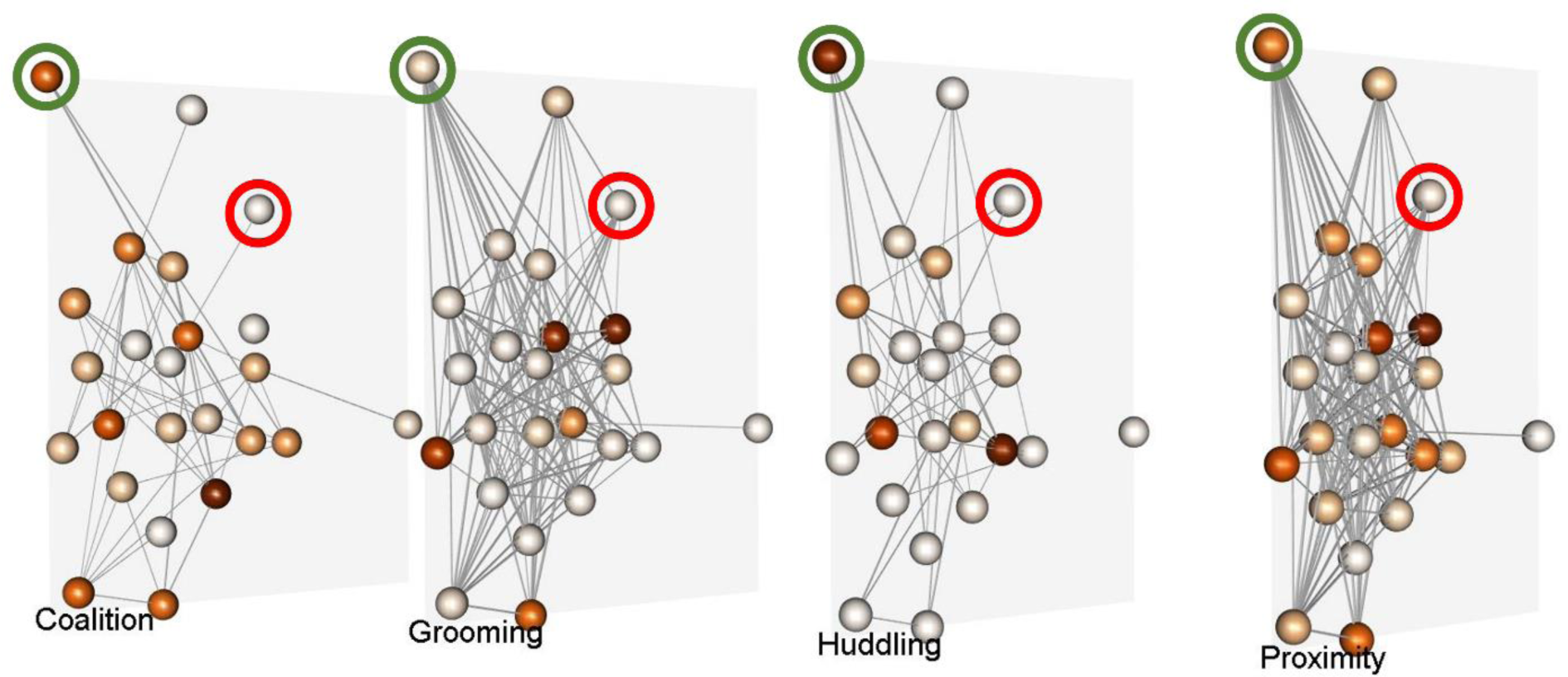
Affiliative multilayer of one of the study groups (HG). Node colour is proportional to eigenvector centrality in each layer with darker nodes being more central, with the same individuals represented in each layer. Edges are rates of interactions of corresponding behaviours. Centrality of some individuals (like the one circled in red) remains the same across all behaviours, while some individuals seem to vary in their centrality across different layers or behaviours (for example, the individual circled in green) indicating that they are more connected (darker) in some behaviours (for example, huddling and coalitionary support) than others (for example, grooming). The affiliative multilayer social networks of the other two study groups (RG and SG) are available in Figure S1 in Supplementary Materials. Made using MuxViz software v2.0.

### Data analyses

To test the socioecological attributes that predicted individual participation in IGC, we used Generalized Linear Mixed Models (GLMM). In each model, the outcome variable was individual counts of participation in IGC. The outcome variable was over-dispersed, so we ran negative binomial models using the ‘glmmADMB’ package [134] in R v4.2.1. As main-effects predictors, we included individuals’ (i) sex, (ii) dominance rank, as well as measures of animals’ social support, specifically their (iii) rate of coalitionary support received, as well as (iv) eigenvector centrality in the coalitionary support network. We also included measures of individual social connectedness, i.e., eigenvector centrality of (v) grooming, and (vi) social proximity networks, as well as their (vii) multilayer affiliative versatility. Since measures of social position (grooming EC, social proximity EC and affiliative eigenvector versatility) were highly correlated with each other (r>0.7[135]), we refrained from including these measures within the same model. To avoid interdependency issues, we also refrained from including the two measures of social support - coalitionary support received and coalitionary support network eigenvector centrality - within the same model. We thus ran six main-effects-only models, and used Akaike Information Criteria (AICc) scores (corrected for small sample size) [136] to select the best-fit. Each of the main-effects only model consisted of (i) sex and (ii) rank, as well as non-interdependent measures of social support (either rate of coalitionary support received or coalitionary support EC) and social connectedness (either grooming EC or social proximity EC or affiliative eigenvector versatility). Based on measures of social/coalitionary support and social connectedness that were selected using the best-fit main-effects only models, we then tested interactions of coalitionary support with rank, sex and single or multilayer measures of social position respectively to investigate if/how participation by individuals with lower socioecological access might be affected by social position (predictions of the underdog hypothesis). Finally, we used AICc to determine the best-fit model(s), assuming that models with lower AICc (change in AIC or DAICc ≥2) explained the data better [136]. In all models, we included Group ID as a random-effects term to account for non-independence of individuals that live in the same group. We set total focal observation time of all our focal animals from all 3 groups as an offset variable. For all models, we checked model diagnostics by examining the raw values, plotting residuals against predicted values, running variance inflation test (VIF) to check for variable collinearity, deviations from normality or obvious outliers. None of them indicated any issues with model validity. For a full candidate model list, refer to Table S3. For all GLMMs we set the critical p value to be 0.05.

## RESULTS

We recorded 478 instances of macaques participating in IGC. We observed inter-individual differences in participation, with 80 out of 116 individuals participating in IGC at least once. The mean individual rate of participation was 0.311± SD 0.37 interactions per hour, with participation ranging from 0 (i.e., some individuals not even participating once) to participating 28 times. (Fig S2).

### Best-fit model selection

Out of the 6 main-effects only models, 2 models fell within a dAICc of less than 2 within each other. Both these models consisted of sex, dominance rank, coalitionary support EC, as well as either social proximity EC or multilayer affiliative versatility. This indicated that coalitionary support EC was a better predictor of participation than rate of coalitionary support received. Moreover, both social proximity EC and affiliative versatility seemed to be equally well-fit, and each better-predictive of IGC participation than grooming EC. Therefore, we decided to proceed with coalitionary support EC (as a measure of social support) and social proximity EC and multilayer affiliative versatility (for measures of social position) to test interaction terms associated with our hypotheses. These analyses identified 2 best-fit models that had very little change in AICc scores (dAICc <0.07) (Table 1).

**Table 1:** Results from two best-fit models.

| *Predictors* | **Model-1 (AICc=528.53)** |  |  | **Model-2 (AICc=528.59)** |  |  |
| --- | --- | --- | --- | --- | --- | --- |
|  | *Estimate* | *Std. Error* | *p-value* | *Estimate* | *Std. Error* | *p-value* |
| (Intercept) | -6.99 | 0.27 | **<0.001** | -6.71 | 0.24 | **<0.001** |
| Dominance rank index | -1.82 | 0.41 | **<0.001** | -1.68 | 0.37 | **<0.001** |
| Sex (male) | 1.32 | 0.22 | **<0.001** | 1.27 | 0.21 | **<0.001** |
| Coalitionary support eigenvector centrality (EC) | 3.63 | 0.67 | <**0.001** | 2.82 | 0.61 | <**0.001** |
| Social proximity eigenvector centrality | 2.40 | 0.71 | **0.001** |  |  |  |
| Coalitionary support EC x social proximity EC | -4.09 | 1.20 | **0.001** |  |  |  |
| Multilayer affiliative versatility |  |  |  | 1.79 | 0.60 | **0.003** |
| Social proximity EC x multilayer affiliative versatility |  |  |  | -2.81 | 1.30 | **0.031** |

### Predictors of individual participation in IGC

The first best-fit model included sex, dominance rank, social proximity EC and coalitionary support EC. It showed a significant negative interaction between coalitionary support EC and social proximity EC (β=-4.09, p=0.001). Exploring this interaction further revealed support for the underdog hypothesis, as individuals with lower social proximity EC participated more in IGC if they had higher coalitionary support EC (Figure 2a). The second best-fit model consisted of sex, dominance rank, multilayer affiliative versatility, coalitionary support EC and a significant negative interaction between coalitionary support EC and multilayer affiliative versatility. Similar to the first best-fit model, for individuals with lower multilayer affiliative versatility, participation in IGC increased with higher coalitionary support than it did for individuals with higher affiliative versatility (β=-2.81, p=0.03) (Figure 2b). Moreover, both best fit models also showed significant effects for sex and dominance rank on participation. More specifically, in support of the Underdog hypothesis, lower-ranking monkeys participated more in IGC than higher-ranking monkeys (β=-1.82, p<0.001 and β=-1.68, p<0.001 respectively) (Figure 3a). On the other hand, in support of the top participant hypothesis, males participated more than females (β=1.32, p<0.001 and β=1.27, p<0.001 for first and second best-fit model respectively) (Figure 3b).

**Figure 2:**
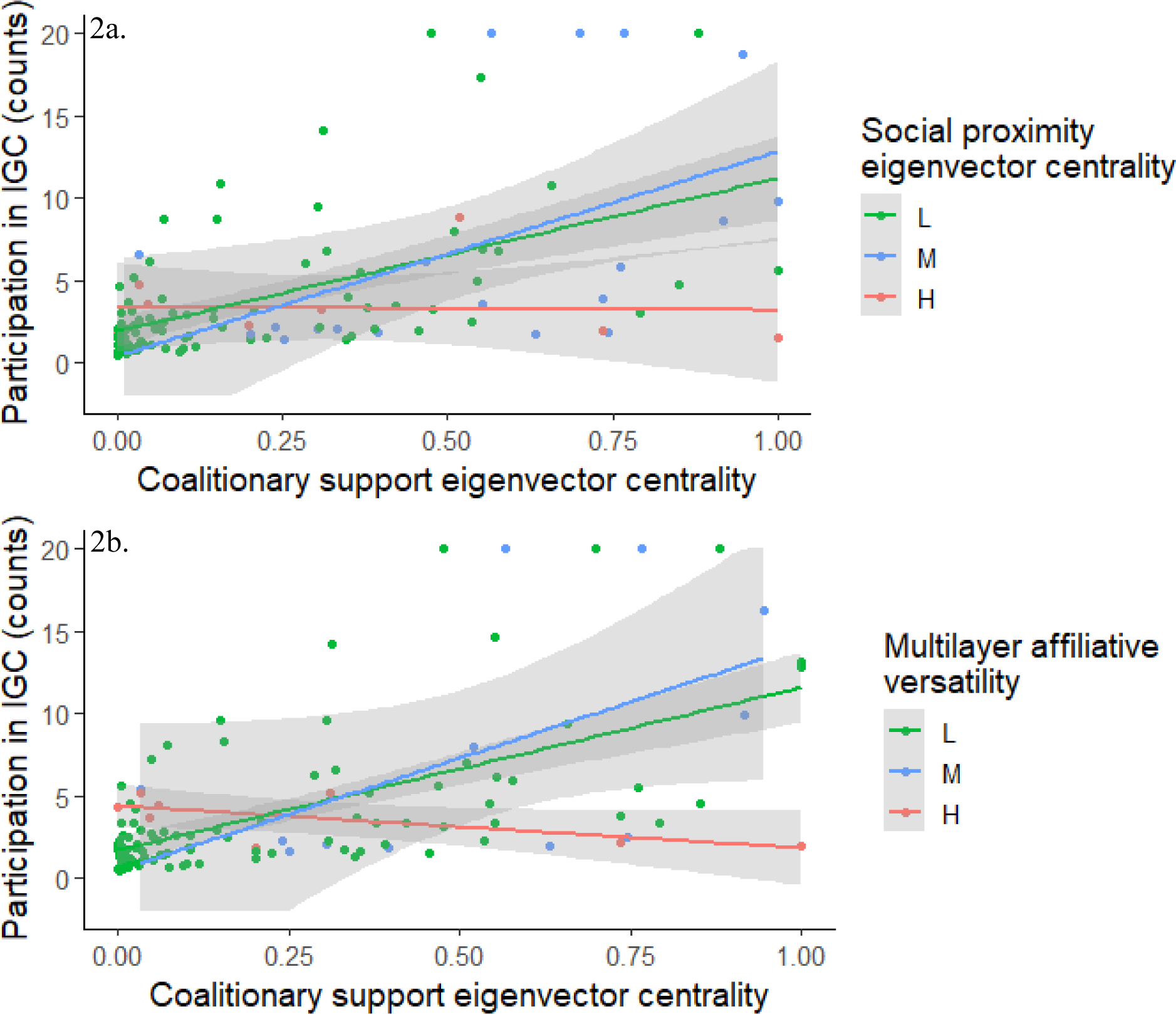
a. **Participation in IGC by coalitionary support eigenvector centrality and social proximity eigenvector centrality** (first best-fit model). b. **Participation in IGC by coalitionary support eigenvector centrality and multilayer affiliative versatility** (second best-fit model). Individuals with lower social proximity EC or multilayer affiliative versatility (in green and blue) are more likely to participate if they have higher coalitionary support EC. Social proximity EC and multilayer affiliative versatility have been converted to a categorical variable for easier visualization. The cut-offs for Low (L), Medium (M) and High (H) are based on natural breaks in social proximity EC/multilayer affiliative versatility values.

**Figure 3:**
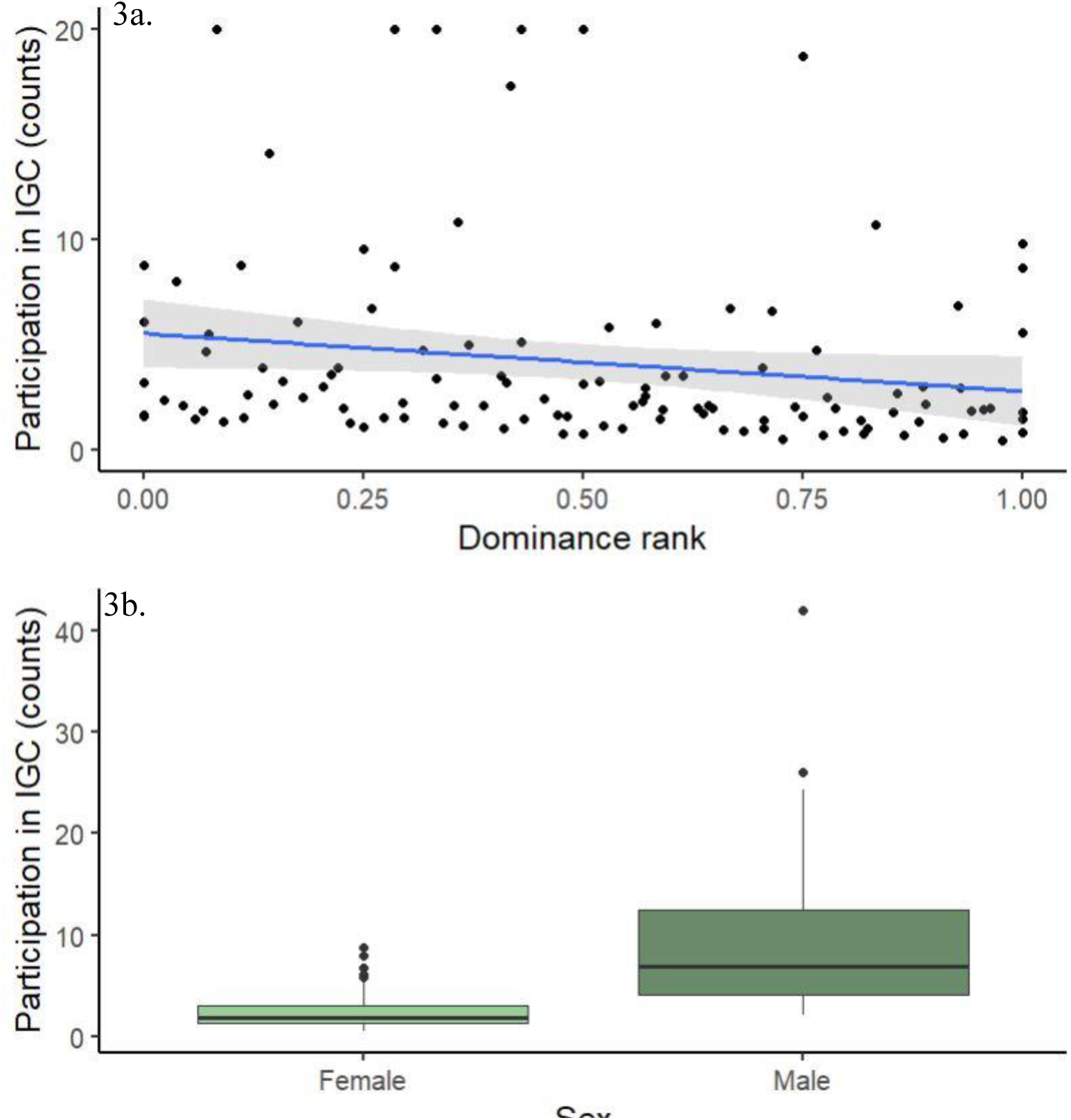
Participation in IGC by a. dominance rank b. sex.

## DISCUSSION

We found strong inter-individual differences driving participation in IGC in urban rhesus macaques, with 69% of the study subjects participating in IGC at least once. Our results revealed that individuals more connected in social support network showed the greatest propensities to participate in IGC. Interestingly, we also found that the effect of coalitionary support on participation was dependent on animals’ social integration in other affiliative networks – support seemed especially important for participation among individuals that were less, compared to more, socially central or integrated within their proximity networks and/or across their multilayer affiliation networks. Moreover, we also found that males and lower-ranking individuals showed more tendencies to participate in IGC than females and high-ranking individuals respectively. Below we discuss these findings in depth, and more generally their evolutionary, health and conservation-related ramifications:

In support of the top participant hypothesis, we found a strong effect of individuals’ intra-group social connectedness and social support on their participation. Surprisingly, the most direct measure of social support (rates of receiving coalitionary support) did not predict participation, but rather participation was better explained by an individual’s overall direct and indirect social support measure (coalitionary support EC). Moreover, social proximity EC and multilayer affiliative centrality were better predictors of participation compared to grooming EC. Greater proximity to conspecifics might encourage animals to take more risks, for example jointly interact with humans [137] or participate in IGC (this study, [6,78]). Moreover, coalitionary support ties within groups may better ‘prime’ animals to participate in IGC, given the links between participation and intragroup cooperation and conflict resolution [55]. Grooming relationships not being as important in the context of IGC could be because grooming may limit the time available for other activities like participation in IGC [89]. Finally, coalitionary support ties seemed more important than rates of receiving coalitionary support. This lack of directional influence of coalitionary support suggests that social integration [138], rather than market-mediated socio-ecological exchange forces [139], may drive participation.

However, we found a significant negative interaction between social proximity EC/multilayer affiliative versatility, and coalitionary support EC predicting participation. Social proximity EC and multilayer affiliative versatility are highly correlated to each other, presumably resulting from individuals occupying socially privileged positions across multiple affiliative behaviours also generally being in proximity to others. It is, therefore, unsurprising that they show similar patterns. In support of the underdog hypothesis, for socially peripheral individuals, having strong coalitionary support ties seems to be a major determinant of their participation decisions since having stronger social support might ameliorate the risks of IGC. On the other hand, it is likely that coalitionary support ties are not nearly as important in terms of governing the participation of individuals that are more socially central, or more socially integrated into, their other affiliative networks. Indeed, participation based on available support to offset the costs of IGC has been seen in other animals like female lions that engage in territorial defence based on the reliability in cooperative tendencies of their companions [140], female vervet monkeys that participated more when males were present [34] and female colobus monkeys (*Colobus vellerosus*) that participated when closely-bonded individuals were present [78]. Here we expand on such findings to reveal how the complex interaction between multiple (rather than single) forms of affiliation can drive animals’ participation in IGC. Frequent IGC is hypothesized to play a major role in the evolution of cooperation and intragroup affiliation, and recently there has been research across the animal kingdom exploring this relationship [51,52,54,55,141]. However, more studies exploring IGC in group-living animals should consider the effect of intragroup social dynamics, and specifically the social integration of individuals within their groups, on IGC to understand the dynamic interplay between the two, and how these influence individual-level decisions to cooperate and compete.

We found an effect of macaques’ dominance rank on their propensities to participate in IGC. Specifically, we found support for the underdog hypothesis rather than the top-participant hypothesis, with low-ranking individuals participating more in IGC than high-ranking individuals. This is a surprising finding, given that most studies on IGC have generally found support for the top-participant hypothesis, i.e. for higher ranking individuals with greater access to resources, and thereby with ‘more to lose’, to participate more in IGC (dogs [32]; meerkats [60]; marmosets, *Callithrix jacchus* [142]; ring-tailed lemur, *Lemur catta* [38]; Japanese macaques, *Macaca fuscata* [64]; bonnet macaques [49]; chacma baboons [143,144]). Yet our results are also consistent with those from other, human-provisioned nonhuman primate populations, wherein participation by individuals with less access to resources (such as lower ranking or indeed immature individuals) has indeed been recorded (e.g., semi-provisioned free-ranging rhesus macaques at Cayo Santiago [63,110,111,145,146]).

There might be several reasons explaining why lower-ranking individuals might participate in IGC in such provisioned populations. First, low-ranking monkeys may have the least access to resources due to intra-group competition and might seek alternate foraging opportunities which might force them to encounter (and engage in) direct competition with extragroup individuals. This might be especially pronounced for human-impacted environments where food is generally more clumped [147,148], resulting in it being more monopolizable by dominant individuals [112]. Moreover, given the high energetic value of anthropogenic food, participation in these habitats might be associated with greater immediate benefits compared to participation in less anthropogenic populations. Second, low-ranking animals are often spatially peripheral within their group [149–151], and might participate in IGC as a result of coming in greater contact with other groups. This might also be driven by greater intergroup contact as a result of higher densities of groups in anthropogenic areas, resulting in these socially and spatially peripheral monkeys encountering extagroup individuals more. Third, subordinate individuals might participate in IGC in the presence of dominant individuals to showcase their cooperative tendencies [42], and thereby gain greater intragroup feeding tolerance [152] or punishment avoidance for defection [55]. Finally, it is conceivable that the effect of rank on participation might be influenced by, or indeed be a byproduct of, other socio-ecological factors such as mating seasonality or specific types of food. For example, increased participation by individuals with higher reproductive access (dominant males) was observed during mating season in Japanese macaques [64].

Finally, in support of the top participant hypothesis, we found that males were more likely to participate in IGC compared to females. This is in line with a large body of literature reporting higher male participation in IGC [73]. In our study groups, males had more access to anthropogenic food than females, which might drive their participation motivations [112]. Moreover, as in many other mammalian taxa, rhesus macaque males are both bigger in size and have less demanding life-history tradeoffs than females [153]. Hence participation may impose fewer costs, and conceivably more benefits, on males compared to females. Another explanation is that in macaques, and in other group-living animals that show similar organizations, males spend more time in the periphery and tend to disperse away from their natal groups, and may therefore face greater intergroup encounters than females [102,154,155] Moreover, limited research on wildlife populations in anthropogenically-impacted environments has to-date revealed that males more so than females tend to engage more frequently with humans (e.g. African elephants, *Loxodonta africana* [156]; macaques: [91,119] and/or jointly aggregate around anthropogenic factors (macaques [137]). Thus, participation by males in IGC might be an artefact of participation in human-wildlife interactions over food, or vice versa, due to which, males may be more likely to encounter members of other groups than females. Interestingly, we did observe female participation in our study group with 50 out of 85 females participating at least once (Figure S4). More generally, other studies have found support for higher female compared to male participation (spotted hyena [72], ring-tailed lemurs [157], vervet monkeys [34], Tibetan macaques, *Macaca thibetana* [97]). Aside from the underdog hypothesis, other explanations for greater female participation pertain to its greater likelihood among species that show female dominance over males (e.g., spotted hyenas [72]) or relaxed dominance relationship between the sexes (e.g., Tibetan macaques [97]), leadership in collective movement behaviour (reviewed in [73]), and/or manipulation of male participation by social incentives (e.g., vervet monkeys [158]). While males may show overall greater frequencies of participation, females may also play important roles in shaping participation decisions.

Overall, our results partially support both of our hypotheses and indicate that participation is not solely based on access to single resource types since individuals with generally more access (males, central individuals) as well as less access (lower-ranking individuals) participated. These findings provide strong evidence that participation in IGC is likely a result of complex decision-making considering various socioecological factors. We uncovered large-scale patterns of participation based on broad proxies of access to socioecological resources, which might give us a snapshot of an individual’s overall balancing of cost and benefits before engaging in these risky behaviours. Future research will use these findings to delve into more proximate, quantitative measures of access to resource quality and quantity, that might interact with animals’ demographic and social status to influence such decision making.

This paper adds to the growing literature applying multilayer social networks to study animal behaviour. Multilayer analysis can help us understand an individual’s role in a group by uncovering how an individual’s actions across various contexts can scale up to group-level consequences such as maintaining group cohesion [100], as well as health outcomes. In fact, a recent study on rhesus macaques found that individuals central in a uniplex (grooming) network had higher measures of inflammation compared to individuals central in a multiplex (grooming and huddling) network [116]. The effect of such seemingly subtle behavioural differences on health outcomes underlies the need to consider such complexities in animal socioecology. In this paper, our finding that a combination of intermediate, scaling up to multilayer measures, were much more likely to influence participation in IGC than animals’ direct social connectedness, clearly highlight the inherent complexity underlying animals’ decision-making, and the utility of such network approaches to disentangle such complexity.

In summary, our study adds to a nascent body of literature that examines how animal socio-ecology in human-impacted or anthropogenic landscapes may impact their propensities to take risks [159], and specifically their participation in IGC. Since IGC constitute a fundamental aspect of the ecology of group-living animals, our findings add to growing perceptions of the need to understand animal ecology in human-impacted environments merely as extensions of ‘natural’ systems with novel, dynamic characteristics, rather than as independent entities [82,160]. More generally, our focus on the behavioural ecology of an urban generalist species that frequently interacts with humans should inform, and indeed lead naturally, to future research in applied ecology, in areas such as ‘conservation behaviour’ [161] and infectious disease ecology [162,163] at these human-wildlife interfaces [164]. Broadly, given that conflict among social groups has been a dominant aspect of human evolution, and continues to be prevalent in modern human societies, understanding individual differences driving such interactions in a species inhabiting similar dynamic habitats can help us take comparative approaches to understand individual motivations that promote or prevent such conflict [165].

## ACKNOWLEDGEMENTS

We thank the Himachal Pradesh Forest Department for their cooperation, and permission to conduct research in Shimla. We are also grateful to Eduardo Saczek, Taniya Gill, Kawaljit Kaur, Benjamin Sipes and Nalina Aiempichitkijkarn for their help in data collection. We also thank Dr. Nitika Sharma for her helpful advice in using MuxViz.

