## Supplementary Materials for "Intragroup sociality drives individual participation in intergroup competition in an urban-dwelling nonhuman primate"

Table S1: **Ethogram for macaque-macaque interactions**
For full ethogram of human-macaque interactions refer to Kaburu et al., 2018 (1)

**Aggression**
**Agg. displacement**: Recipient is displaced from where they are by another individual.

**Agg. threat**: initiator stares with a widely open mouth and/or flashing eyebrows – staring with exaggerated raising and then release of the eyebrows, and/or ear flap-staring while quickly pulling the ears forward and then releasing them.

**Agg. lunge**: Makes rapid quick leaps towards the recipient.

**Agg. slap**: Hitting or striking another individual.

**Agg. grab**: To take hold of another monkey, suddenly and roughly.

**Agg. chase**: the aggressor runs threateningly after the recipient.

**Agg. bite**: physical contact involving teeth.

**Agg. wrestle**: Prolonged physical contact with another individual where one animal attempts to restrain or restrict the other's movement (e.g., gripping and struggling).

**Agg. pin**: Holding the recipient for at least 3 seconds.

**Agg. push**: Recipient is pushed by the initiator's limbs and/or body.

**Agg. redirection**: Aggressive behavior or displacement by one monkey toward another monkey in response to an approach or aggression by a third monkey.

**Impartial intervention**: The intervener is not functionally supporting one animal/party over the other, but attempts to stop the conflict, either by walking between the two animals/parties, or threatening both parties.

**Aggressive display**: Shaking or rocking a tree limb or other structure (e.g., man-made structure) to generate noise and attract the attention of conspecifics.

**Submission**
**Silent bared teeth (SBT):** Lips are pulled back in an exaggerated manner exposing teeth (also called fear grimace). Animals' brows are pushed together in a way that makes the animal appear frightened; may or may not be accompanied by a scream. If followed by scream, score SC only.

**Turn away:** Moving the upper body away while not changing location

**Avoiding:** Walking or jogging (i.e. running not at full speed) out of arm's reach of the aggressor

**Flee:** Running away from another animal following aggression

**Scream:** Loud, shrill vocalization

**Recruitment**: Solicitation of a third party, not currently involved in the conflict, while engaged in antagonistic behavior directed at their opponent. Must involve one or more of the following:

rump present, looking back and forth between the third party and the opponent, or approach while continuing to threaten the third monkey.

**Affiliation**

**Grooming**: An individual cleaning or manipulating the fur of another individual

**Huddling**: An individual is ventrally oriented towards another individual for at least 5 seconds

**Coalitionary support**: An individual supports one individual/party against another individual in aggressive interactions.

**Lip-smacking**: Rapid movement of the pursed lips, up and down, can occur in conjunction to or preceding grooming, huddling and mating.

**Play:** Non-aggressive chasing, bouncing, tumbling, grabbing, wrestling, soliciting, and mock biting of another monkey.  These behaviors are often seen with a relaxed open mouth ‘play face’.

**Neutral**

**Non-sexual mounting**: A monkey grabs the hind legs of another monkey with his/her own hind feet and places his/her hands on the lower back of the recipient, thus hoisting himself/herself off of the ground; may include thrusting but looks sloppy and is shorter in duration compared to a sexual mount.

**Non-sexual present**: A monkey presents his/her rump to an individual of the same sex. Is often followed by an affiliative behavior.

**Neutral grab**: A monkey holds on to a monkey of the same sex; can be during aggressive or affiliative contexts.

**Sexual**

**Sexual mounting**: A monkey grabs the hind legs of another monkey with his/her own feet and places his/her hands on the lower back of the recipient, thus hoisting himself/herself off the ground; must include deliberate thrusting; the recipient often looks back, lipsmacks, or grabs the mounter; may be accompanied by screams; the mounter may huddle and hold on to the recipient after the mount.

**Sexual presenting**: An animal offers his/her rump to another.  May include an upright tail, arched back, or touching head to the ground with straight legs.  Includes sexual rump presents.

**Sexual inspection**: Involves a male sniffing or touching a female’s genitalia.

Table S2: **Variable correlations:**

| | Dominance rank index | Rate of receiving coalitionary support | Coalitionary support eigenvector centrality (EC) | Grooming EC | Social proximity EC | Multilayer affiliative versatility |
|---|---|---|---|---|---|---|
| Dominance rank index | 1.0000 | 0.2006 | 0.3343 | 0.2507 | 0.5593 | 0.3159 |
| Rate of receiving coalitionary support | 0.2006 | 1.0000 | 0.2046 | 0.0967 | 0.1991 | 0.0378 |
| Coalitionary support eigenvector centrality (EC) | 0.3343 | 0.2046 | 1.0000 | 0.1772 | 0.4034 | 0.2812 |
| Grooming EC | 0.2507 | 0.0967 | 0.1772 | 1.0000 | 0.6579 | 0.8680 |
| Social proximity EC | 0.5593 | 0.1991 | 0.4034 | 0.6579 | 1.0000 | 0.7836 |
| Multilayer affiliative versatility | 0.3159 | 0.0378 | 0.2812 | 0.8680 | 0.7836 | 1.0000 |

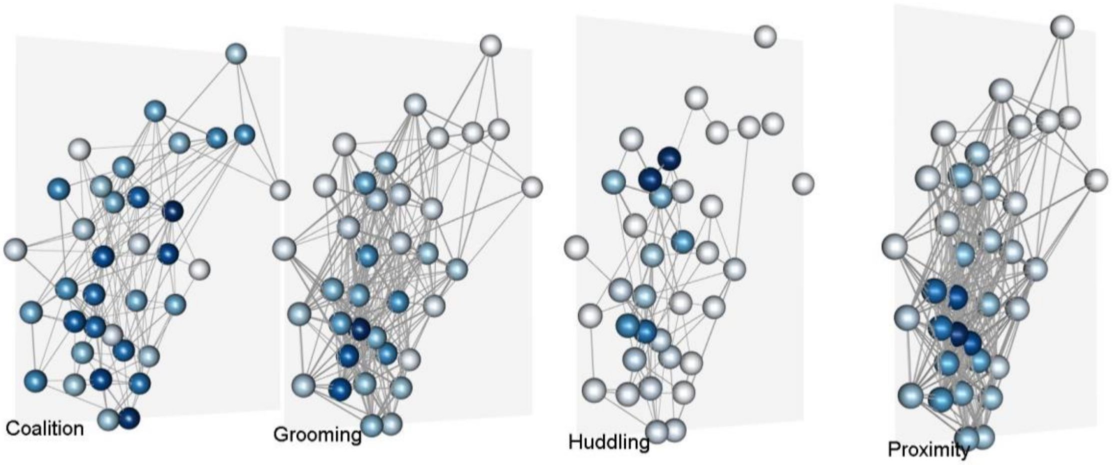

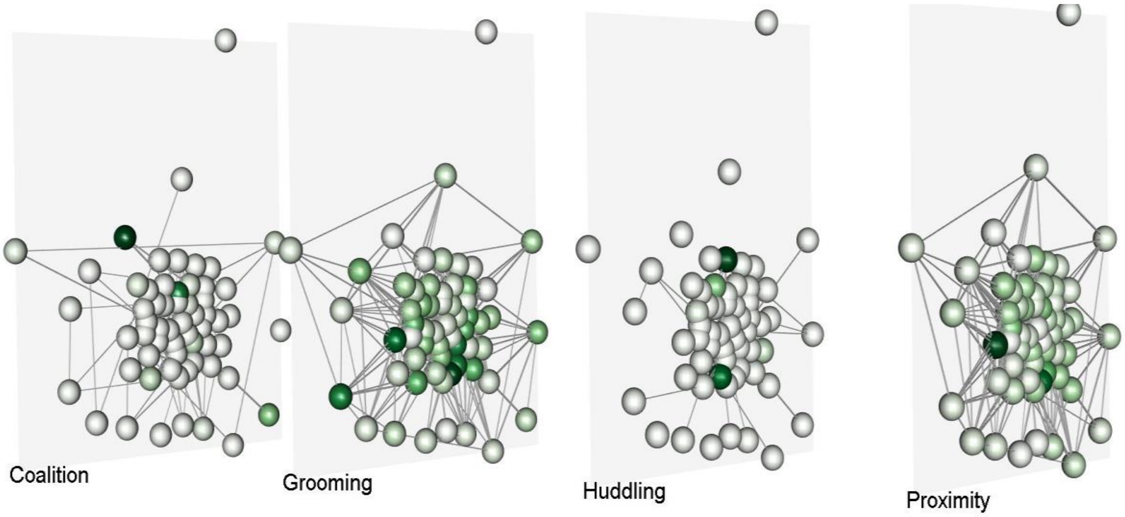


Figure S1: **Affiliative multilayer social network of two of the study groups**: RG (top), SG (bottom).

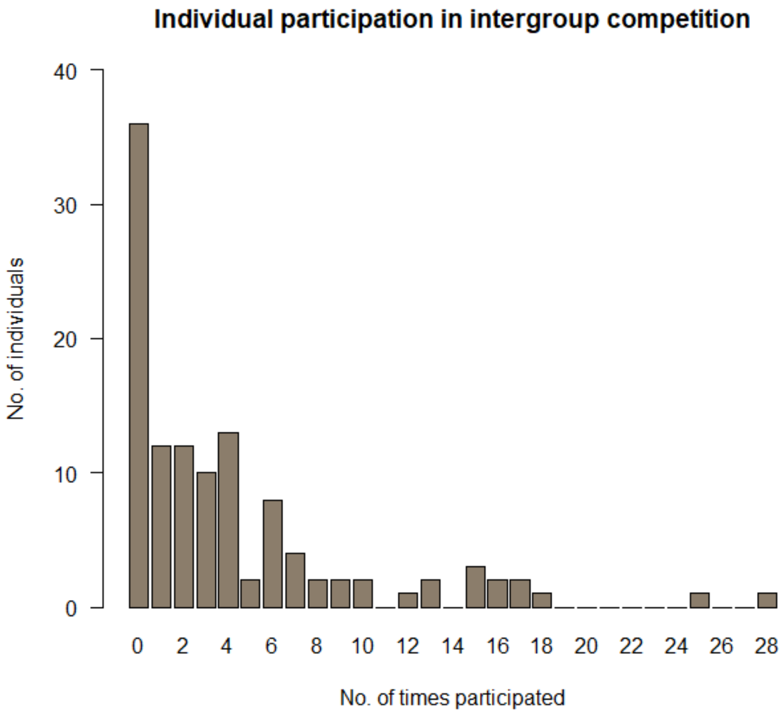


Figure S2: Individual participation in intergroup competition

Table S3: **Full candidate model list**.

Model abbreviations: *RankIndex*: standardized dominance rank, *CSRec*: rate of receiving coalitionary support, *Gr*: grooming eigenvector centrality, *CS*: coalitionary support eigenvector centrality, *Prox*: social proximity eigenvector centrality, *Aff_Ver*: multilayer affiliative versatility. Final best-fit models in bold.

| No. | Model | AICc |
|---|---|---|
| 0 | Null | 569.912 |
| 1 | Participation~RankIndex+Sex+CSRec+Gr | 541.893 |
| 2 | Participation~RankIndex+Sex+CS+Gr | 532.727 |
| 3 | Participation~RankIndex+Sex+CSRec+Prox | 537.869 |
| 4 | Participation~RankIndex+Sex+CS+Prox | 530.277 |
| 5 | Participation~RankIndex+Sex+CSRec+Aff_Ver | 539.057 |
| 6 | Participation~RankIndex+Sex+CS+Aff_Ver | 530.713 |
| 7 | Participation~RankIndex*CS+Sex+Prox | 531.044 |
| 8 | Participation~RankIndex+CS*Sex+Prox | 532.532 |
| 9 | Participation~RankIndex+Sex+CS*Prox | **528.528** |
| 10 | Participation~RankIndex*CS+Sex+Aff_Ver | 531.602 |
| 11 | Participation~RankIndex+Sex*CS+Aff_Ver | 532.946 |
| 12 | Participation~RankIndex+Sex+CS*Aff_Ver | **528.592** |

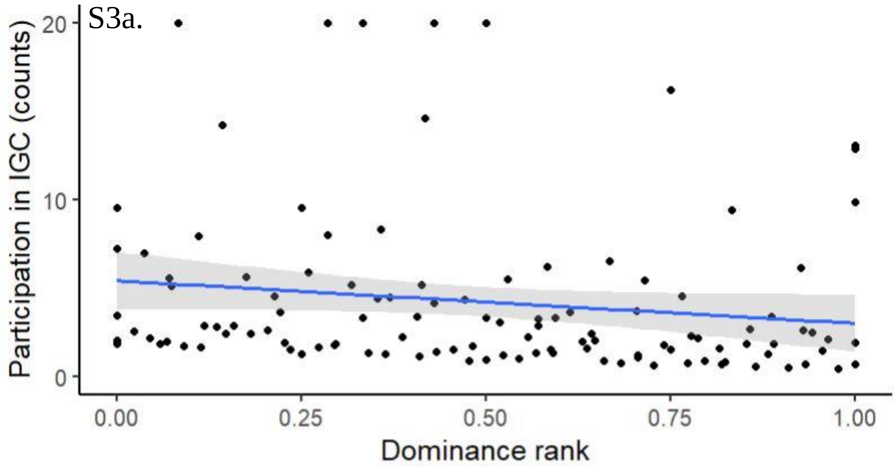


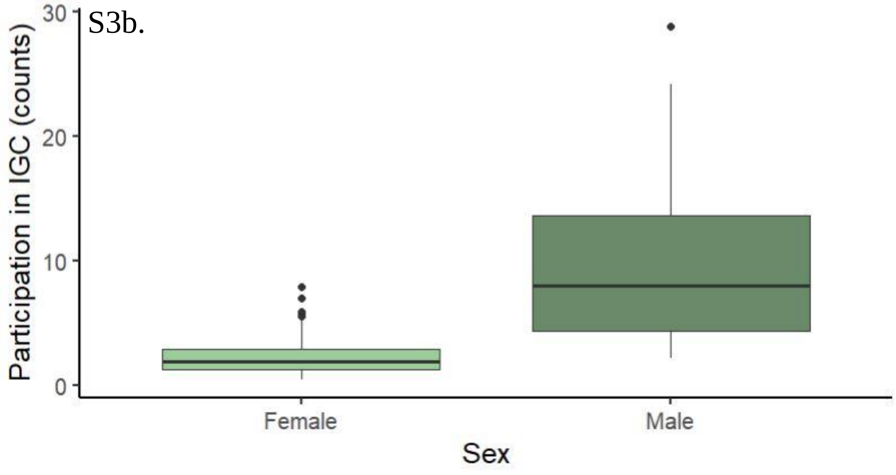

Figure S3: **Participation in IGC by a. dominance rank and b. sex** (second best-fit model)

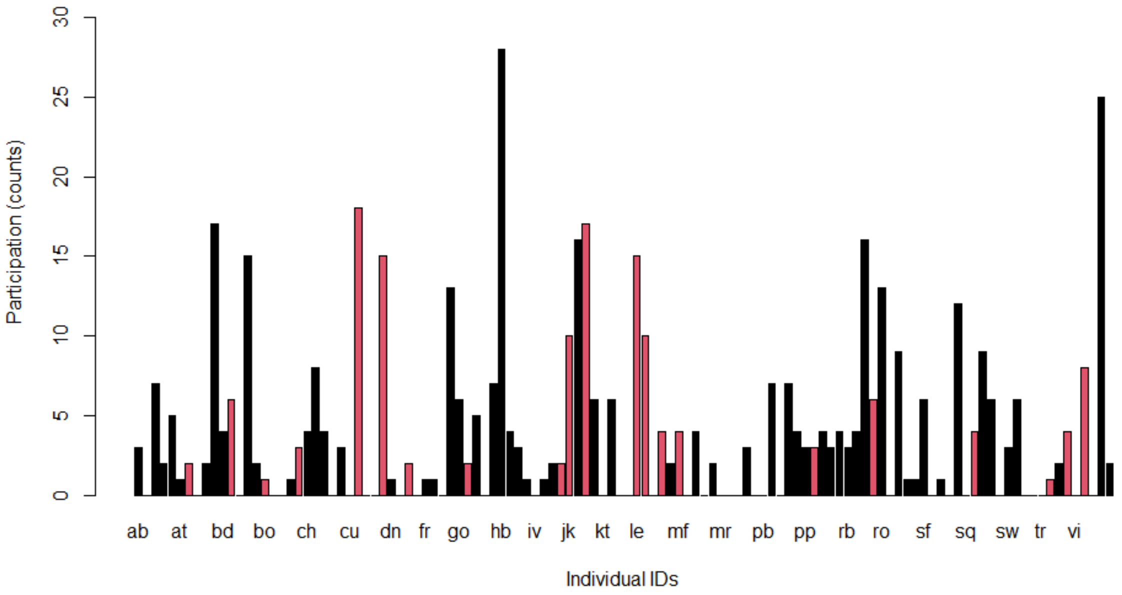


Figure S4: **Differential participation by male and female rhesus macaques**. Black: males, red: females (N=116, males=31, females=85)
